# Mitochondrial perturbation adapts the proteome from early to advanced immune responses

**DOI:** 10.64898/2026.09.07.749965

**Authors:** Jessica Alvarez, Sofia Rivera, Sarahbeth Roberts, Natalia Herrera, Tyron Chang, Krishna Kanth Chitta, Jeon Lee, Andrew Lemoff, Dustin C. Hancks

## Abstract

Dysregulated immunity, a hallmark of many human diseases, co-occurs with mitochondrial dysfunction and is commonly associated with misprimed primary immune signaling. While transcriptionally well-characterized, the impact of mitochondria on the host response at the protein level is less clear. Using *in vitro* and *in vivo* approaches including proteotranscriptomics, our data suggest that OXPHOS promotes expression of early, cell autonomous immune proteins whereas mitochondrial perturbation favors mediators of cell extrinsic responses like inflammation. This response is independent of immune cues, time-dependent, conserved, and occurs across tissues in mouse models of mitochondrial dysfunction. These data illustrate unappreciated roles for mitochondrial state in adapting host responses at the protein level, which have implications for complex disease etiology and the ancestral origins for eukaryotic immune sensing.

## Main Text

Dysregulated mitochondrial activity is a universal hallmark for many human diseases that co-occurs with sterile inflammation in chronic illnesses (*1, 2*). Well-known diseases with mitochondrial dysfunction are type II diabetes, obesity, cancer, Parkinson’s, Alzheimer’s, and autoimmunity (*3*). While mitochondrial dysfunction is historically related to bioenergetics, new insights have expanded its definition to include the increasing diversity of mitochondrial activities (*4*). Given extensive crosstalk between the organelle and the cell involving the electron transport chain (ETC), redox functions, TCA cycle, and lipids, perturbation of mitochondria disrupts homeostatic metabolism.

Beyond core homeostatic activities like oxidative phosphorylation (OXPHOS) (*5*), mitochondria are vital orchestrators of host defense against pathogens (*6, 7*). For instance, mitochondria trigger cell intrinsic host responses by inducing type I interferon (IFN) transcription and NF-kB signaling via RLRs/MAVS (*8*). Mitochondrial perturbations also cue the progression of the host response. Two appreciated examples are cytosolic release of both cytochrome c (*9*), which advances apoptosis, and mtDNA (*10*), which activates both type I IFN transcription and the NLRP3 inflammasome. Downstream of cell autonomous defenses, mitochondria shape immune cell effector functions over both short timeframes, like the production of immunomodulatory metabolites by macrophages (*11*), and longer timeframes through trained immunity (*12*). Broadly, shifts in metabolic programs are associated with effector states in immune cells (*13, 14*). This is illustrated by cytokine and pathogen-associated molecular pattern (PAMP) activation of immune cells. For example, activated pro-inflammatory macrophages are typically glycolytic (*15*). In contrast, anti-inflammatory macrophages often favor an OXPHOS state (*16*). Cell-intrinsically, mitochondrial rewiring is thought to function by regulating the magnitude of the host response. Many diseases with mitochondrial dysfunction, an extreme type of mitochondrial rewiring, are comorbidities for infections such as influenza A (*17*). The frequent intersection of mitochondrial functions and host immunity is credited to a “break in endosymbiosis” (*1*). Thus, properly regulated mitochondrial activity is vital for an effective host response and overlaps with various facets of immune defense.

Mitochondrial rewiring is a signature of immune and non-immune cells infected by diverse intracellular pathogens (*14, 18*). In infected cells, mitochondrial rewiring is attributed to a combination of damage, biosynthetic demands of pathogen replication, and host survival. Although glycolysis is a metabolic program commonly upregulated during mitochondrial dysfunction, glucose-rich cell culture media, which promotes glycolysis, is used in majority of host defense studies. In contrast, galactose or low glucose media, which favors OXPHOS and mimic the cellular state similar to what a pathogen first encounters, is rarely used. Media-mediated rewiring of mitochondrial activity has identified proteins that regulate OXPHOS (*19*), cell death (*20*), and poxvirus infection (*21*). As mitochondrial activities are extensively intertwined with cellular processes, the consequences of their disruption are likely far-reaching. Ill-defined mito-immune functions are also implied by recently identified ultraconserved ETC factors that are regulated by immune cues and encoded by diverse viruses (*22*). To date, the impact of mitochondria on the proteome is largely unclear and understudied compared to its role in transcriptional induction of immune responses. Our study finds that mitochondrial activity imparts a conserved logic to the proteome, but not transcriptome, that demarcates expression into two classes – an early, cell autonomous response and an advanced response, which involves inflammatory effectors.

## Results

### Mitochondrial reprogramming stratifies expression of proteins with immune activities

While mitochondrial reprogramming occurs in non-immune cells during infection, consequences of this altered state are less studied compared to immune cells. As a reductive model, we leveraged uninfected cells treated with immune signals to study non-antagonized host responses in culture. This model may also inform activities in bystander cells, a key infection determinant proximal to initially infected cells that the infection spreads into. Using our published protocol (*21*) adapted from the mitochondrial field, we rewired mitochondrial activity through media supplementation to promote OXPHOS or glycolysis in human A549 lung epithelial cells. This line was selected for its widespread use in infection studies and its many intact host defense pathways (*21–23*).

Briefly, cells were grown in media either supplemented with galactose to favor OXPHOS or glucose to promote glycolysis (Fig.1A). Next, cells were either left untreated or treated with type I (IFN-α) or type II IFN (IFN-γ) to trigger antiviral states (Fig.1B). Both IFNs have pleiotropic effects on immune and non-immune cells that influence infection and disease outcomes (*24*). IFNs act, in part, by inducing hundreds of mRNAs encoding immune factors. Consistent with metabolic reprogramming (Fig. 1C), A549 cells in galactose-media, with or without IFNs, displayed increased mitochondrial respiration relative to glycolysis based on the increased ratio of oxygen consumption rate (OCR) to extracellular acidification rate (ECAR). Conversely, cells in glucose-media displayed a lower OCR/ECAR ratio.

**Fig. 1.**
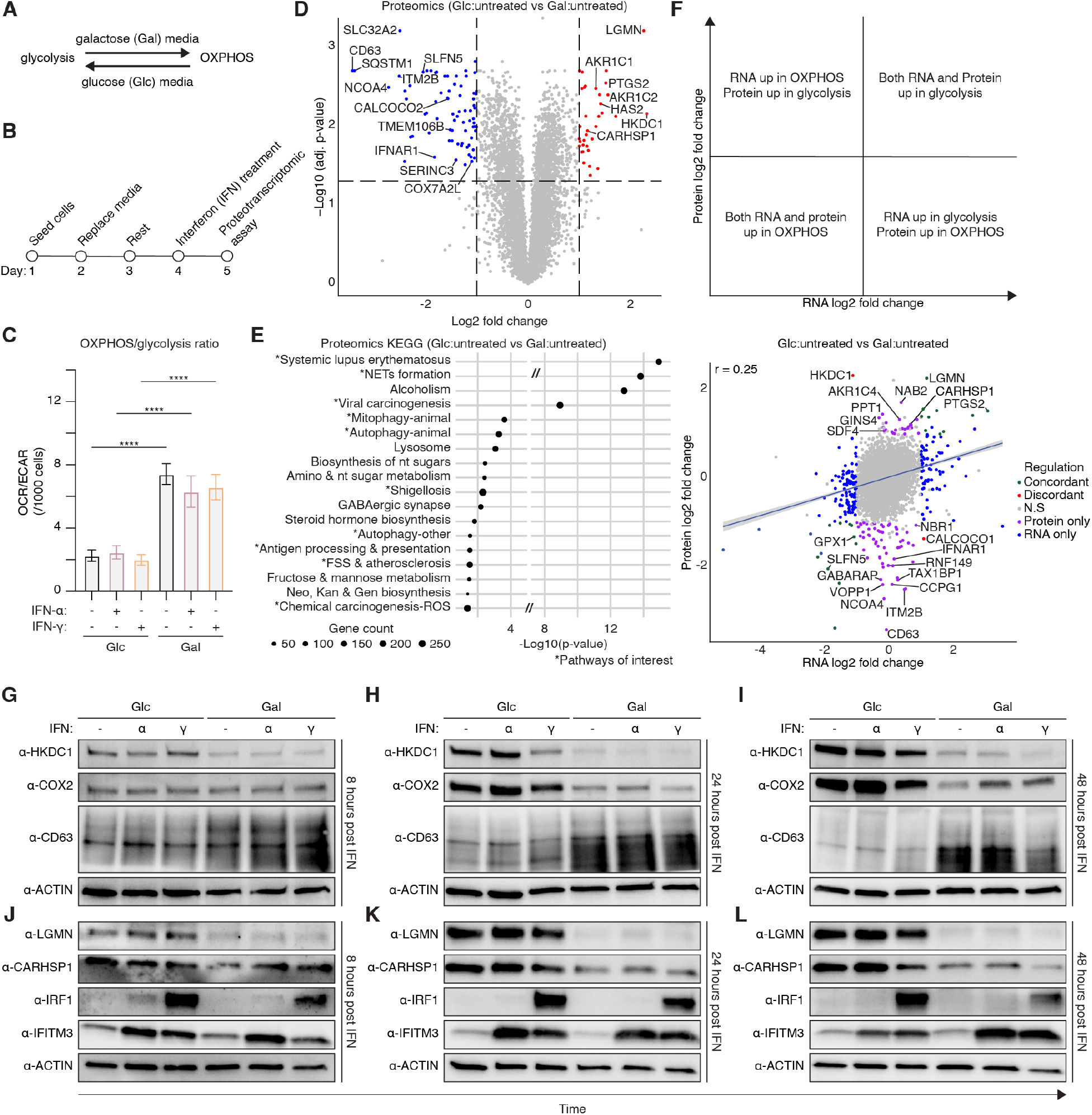
Mitochondrial rewiring stratifies the host response proteome. (**A**) Metabolic rewiring using glucose (Glc) and galactose (Gal) culture conditions in human A549 lung epithelial cells. (**B**) Experimental timeline. (**C**) Basal ratio of oxygen consumption rate (OCR) to extracellular acidification rate (ECAR) using Seahorse analysis; unpaired t-test **** p ≤ 0.0001. (**D**) Differential protein abundance comparing Glc:untreated vs. Gal:untreated. Significantly regulated proteins are highlighted in OXPHOS (blue), glycolysis (red), and non-significant proteins (grey). Proteomics data: adj. p < 0.05; fold-change > 1; greater than 1 peptide. (**E**) KEGG pathways for Glc:untreated vs. Gal:untreated. “*” Pathways of interest. (**F**) Proteotranscriptomic correlation plots comparing Glc:untreated vs. Gal:untreated. Scatter plots show RNA log2 fold changes vs. protein log2 fold changes. Genes are color-coded by regulation status: concordant (green), discordant (red), protein-only (purple), RNA-only (blue), and non-significant (grey). The blue line indicates linear regression with 95% confidence interval. (**G-L**) Protein samples from the indicated time points (hours: 8, 24, 48) were harvested and analyzed for expression by western blot. Cells in Glc or Gal supplemented media were treated with either IFN-α or IFN-γ. Conditions are Glc (25 mM; standard media), Gal (10 mM), and IFN concentration 1000 units/mL.

To test for effects of mitochondrial reprogramming on protein levels, we carried out quantitative proteomics using cells grown in either glucose or galactose media with no IFN-treatment (Fig. 1, D to F, Table S1, Data S1). As expected, metabolic proteins like the hexokinase HKDC1 (*25*) and COX7A2L (*26*), an ETC accessory subunit, were directionally regulated between media (Fig.1D). Unexpectedly, numerous immune proteins and pathways were also differentially expressed between media. Notably, galactose media favored expression of proteins with roles in the early stages of antiviral defense (IFNAR1, SERINC3 (*27*)), negative regulation of immunity (LRP10, APOE), and viral entry (TMEM106B (*28*), CD63 (*29*)). Also increased in galactose media were several autophagy receptors, which have immunoregulatory activities (SQSTM1/p62 (*30*), NCOA4 (*31*), TAX1BP1 (*32, 33*). In contrast, glucose media increased inflammatory regulators: COX2 (also known as PTGS2), LGMN (also known as AEP, δ-secretase), and the TNF-α positive regulator CARHSP1 (*34*). Surprisingly, glucose versus galactose proteome differences showed enrichment for several KEGG immune pathways (systemic lupus erythematosus, neutrophil extracellular trap formation, viral carcinogenesis, and shigellosis) (Fig.1E). Also enriched were KEGG pathways that involve conserved cellular functions, which bridge metabolism and cell autonomous defenses (autophagy, mitophagy and lysosome). A comparable immune signature pattern was also evident by GO analysis (Fig. S1A). To test whether mitochondrial rewiring was acting on the RNA, protein level, or both, we integrated the proteomics data with matching RNA-seq data (Fig.1F, Table S1, Data S2). Proteotranscriptomics analysis highlighted that many of the proteome differences between glucose and galactose media, which included infection determinants (IFNAR1, CD63, SLFN5, MFGE8), occurred only at the protein and not RNA level (correlation 0.25, p-value 1 x 10^-84^). Additionally, majority of pathways enriched at the protein level for glucose compared to galactose media were not enriched in the RNA-seq (Fig. S1, B and C). These data suggest that metabolic reprogramming alone is sufficient to impart an immune signature on the proteome that is not evident in the transcriptome.

### Differential expression of immune proteins by metabolic state is time dependent

Temporal regulation of host responses by immune cues is essential, and dysregulation of this timing can lead to disease. To test whether regulation of immune proteins by mitochondrial reprogramming occurs over time, we carried out a time-course experiment including IFN-treatments. In agreement with reprogramming, the hexokinase HKDC1 (*25*) was elevated early in glucose media and differences became starker over time (Fig.1, G to I). Similarly, CARHSP1, LGMN, and COX2, which have immunomodulatory roles, were generally increased in glucose media with or without IFN-treatments (Fig.1, G to L). In contrast, CD63, a known viral attachment factor (*29*), was upregulated starting at 24 hours and sustained in galactose media. As controls, we included two factors we identified previously regulated at the protein level by mitochondrial reprogramming (*21*), the pro-inflammatory transcription factor IRF1 and the early antiviral protein IFITM3. Over time, IRF1 and IFITM3 became more divergently expressed based on media (Fig.1, J to L). At 48 hours post-treatment, IRF1 was elevated in glucose:IFN-γ whereas IFITM3 was increased in galactose IFN-α and IFN-γ-treated cells. These data indicate there is a time-component to differential and directional regulation of immune proteins by mitochondrial rewiring not reliant on interferons.

### Metabolic state influences expression of proteins with immune functions in IFN-treated cells

Given that the interferon-stimulated genes (ISGs) IRF1 and IFITM3 are regulated by metabolic state, we next asked whether other factors in IFN-treated cells were differentially expressed by media-mediated mitochondrial rewiring. With our cutoff of > 1 peptide, IRF1 and IFITM3, which generated one peptide, were not included. Nevertheless, classic ISG proteins and immune pathways were induced by IFNs relative to untreated, as expected, in both glucose and galactose media [IFN-α: IFITs, OAS, and TRIMs (Fig. S2, A and B, Fig. S3, A to D); IFN-γ: GBPs, HLA, and TAPs (Fig. S2, C and D, Fig.3, E to H)].

Comparing across media and IFN-treatments, principal component analysis (PCA) analysis indicated that both proteomes (Fig. S4A) and transcriptomes (Fig. S4B) for glucose:IFN-α and galactose:IFN-α treated cells were more similar to glucose:untreated and galactose:untreated, respectively, than they were to each other. Conversely, proteomes and transcriptomes for glucose:IFN-γ and galactose:IFN-γ treated cells were more related to each other than any other condition. For the IFN-α treated cells, glucose and galactose media resulted in 91 differentially expressed proteins (Fig.2A, Table S1). In contrast, 23 proteins were differentially expressed in IFN-γ treated cells due to media (Fig.2B, Table S1). Congruent with mitochondrial rewiring, metabolic pathways such as OXPHOS (IFN-α)(Fig.2C, Fig. S5A) and fatty acid elongation (IFN-γ)(Fig.2D, Fig. S5B) were enriched for glucose:IFN-α vs. galactose:IFN-α as well as glucose:IFN-γ vs. galactose:IFN-γ, respectively. Like glucose:untreated vs. galactose:untreated (Fig. 1E), mitophagy, autophagy, and lysosome KEGG pathways were also enriched in proteome differences linked to media for both IFN-α (Fig. 2C) and IFN-γ (Fig. 2D).

**Fig. 2.**
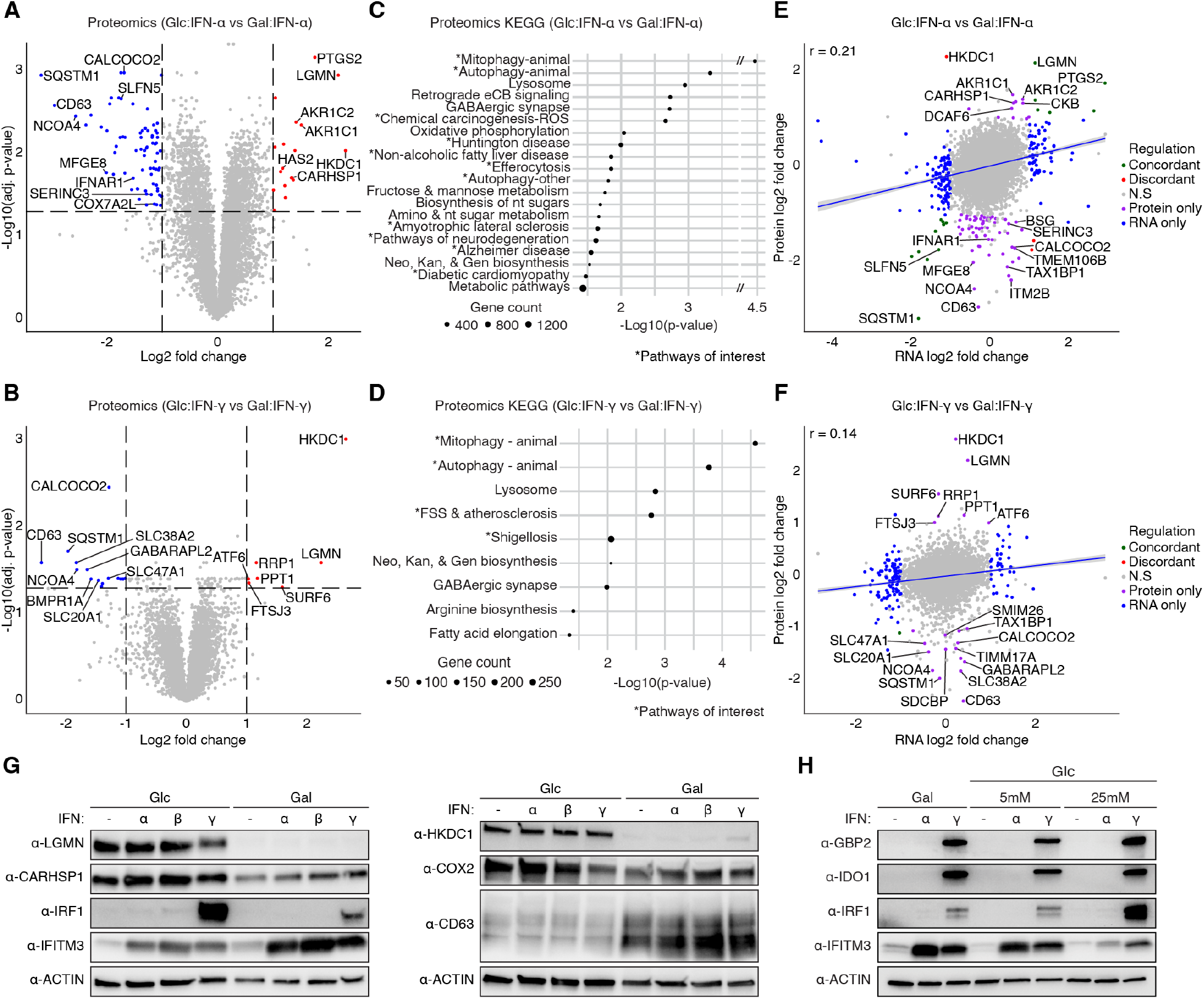
Mitochondrial rewiring impacts expression of numerous immune response proteins in IFN-treated cells. (**A, B**) Differential protein abundance comparing Glc:IFN-α vs. Gal:IFN-α and Glc:IFN-γ vs. Gal:IFN-γ, respectively. Significantly regulated proteins are highlighted in OXPHOS (blue), glycolysis (red), and non-significant proteins (grey). Proteomics data: adj. p < 0.05; fold-change > 1; greater than 1 peptide. (**C, D**) KEGG pathways for Glc:IFN-α vs. Gal:IFN-α and Glc:IFN-γ vs. Gal:IFN-γ, respectively. (**E, F**) Proteotranscriptomic correlation plots comparing Glc:IFN-α vs. Gal:IFN-α and Glc:IFN-γ vs. Gal:IFN-γ, respectively. Scatter plots show RNA log2 fold changes vs. protein log2 fold changes. Genes are color-coded by regulation status: concordant (green), discordant (red), protein-only (purple), RNA-only (blue), and non-significant (grey). The blue line indicates linear regression with 95% confidence interval. (**G**) Western blot for ISGs using protein lysates from human A549 cells grown in Glc/Gal media treated with IFNs. (**H**) Western blot of ISGs for IFN-treated cells in Gal (10mM) or Glc (5mM, 25mM) media. Glc: glucose, Gal: Galactose. Conditions are Glc (25 mM; standard media), Gal (10 mM), and IFN concentration 1000 units/mL.

Although host response factors such as SLFN5, IFNAR1, and COX2 were differentially expressed between glucose:IFN-α and galactose:IFN-α, there was no clear enrichment for host defense pathways (Fig. 2, A and C) comparable for IFN-α relative to untreated (Fig. S3, A to D). Instead, IFN-α treatment between glucose and galactose showed KEGG pathway enrichment for diverse diseases, which display mitochondrial dysfunction (*3*) (Fig. 2C), that was not reflected in the RNA-seq data (Fig. S6, A and B). For proteome differences between glucose:IFN-α and galactose:IFN-α, disease pathways enriched were Huntington’s disease, nonalcoholic fatty liver disease, ALS, pathways of neurodegeneration, Alzheimer’s disease, and diabetic cardiomyopathy. A complementary analysis that incorporated IFN-induction highlighted a similar amount of protein differences (Table S1) and similar pathway enrichment for glucose:IFN-α relative to galactose:IFN-α (106 proteins) (Fig. S7). However, this analysis unmasked greater differences for glucose:IFN-γ vs. galactose:IFN-γ (55 proteins) (Fig. S7B, Table S1).

Next, we assessed how many of these differences in IFN-proteomes were happening at either the RNA or protein level. As anticipated, we found that for each media, IFN-α and IFN-γ treated RNA-seq relative to untreated showed enrichment for antiviral pathways (Fig. S8 and Fig. S9), with a strong correlation for the RNA and protein differences (Fig.S10, A to D, Table S1). However, similar to glucose:untreated and galactose:untreated, many of the proteome changes between media for IFN-treated cells did not correlate well with differences at the RNA level (Fig.2, E and F, Table S1) [IFN-α:glucose vs. IFN-α:galactose, (r = 0.21); IFN-γ:glucose vs. IFN-γ:galactose (r = 0.14)]. This lack of correlation was corroborated by the second analysis incorporating IFN-induction (Fig.S11 and S12). Underscoring the breadth of this regulation, cells treated with IFN-β, another type I IFN, also displayed galactose upregulation of IFITM3 (Fig. 2G). This metabolic effect was also not limited to galactose as evidenced by the differential expression of IRF1 and IFTIM3 in low-glucose media, a complementary means to promote OXPHOS (Fig.2H). These data demonstrate that the metabolic state has a marked and directional effect on immune protein levels in IFN-treated cells.

### Regulation of immune proteins by metabolic state is evolutionary conserved

To test whether metabolic regulation of immune proteins was conserved in immune cells, similar experiments were performed with mouse BV2 microglial cells. BV2 cells are commonly used in viral studies (*35, 36*). As expected, BV2 cells in galactose media displayed upregulation of respiration relative to glycolysis compared to cells in glucose media (Fig.3A). Following IFN-treatments, BV2 cells also upregulated *Irf1* and *Ifitm3* RNA, but not *Lgmn* (Fig.3B). We found that IRF1 protein was elevated in glucose conditions across time points whereas IFITM3 was elevated early in glucose and, like A549 cells, increased in galactose at the late timepoint (Fig.3, C to E). Consistently, we previously observed glucose-mediated IRF1 upregulation in IFN-γ treated mouse embryonic fibroblasts and a cat kidney line (*21*). Likewise, LGMN protein levels became markedly different over time with increased expression in glucose media (Fig.3, C to E). These data suggest that temporal and directional regulation of select immune proteins by mitochondrial reprogramming is conserved in different cell types and other species.

**Fig. 3.**
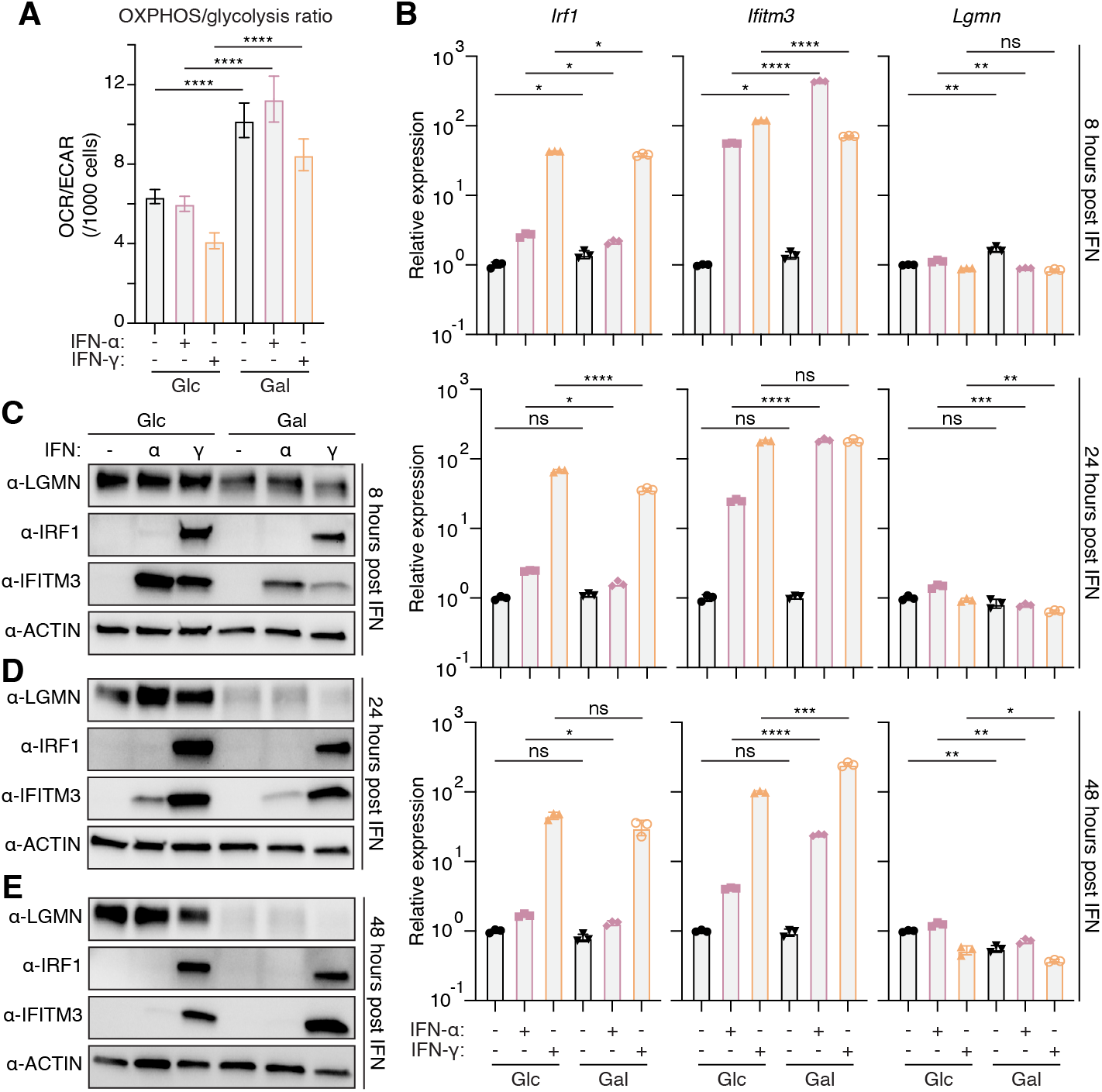
Mitochondrial-mediated adaptation of the host response proteome is evolutionarily conserved. Mouse BV2 microglial cell samples from the indicated time points (hours: 8, 24, 48) were harvested and analyzed for RNA and protein expression. (**A**) Basal ratio of oxygen consumption rate (OCR) to extracellular acidification rate (ECAR) using Seahorse analysis. (**B**) qPCR of immune factors from BV2 cells in Glc or Gal supplemented media treated with either IFN-α or IFN-γ over 8, 24, and 48 hours, respectively. (**C-E**) BV2 western blots of immune proteins from cells in Glc or Gal supplemented media treated with either IFN-α or IFN-γ over 8, 24, and 48 hours, respectively. Glucose (Glc) and galactose (Gal). Conditions are Glc (25 mM; standard media), Gal (10 mM), and IFN concentration 100 units/mL. Statistical analysis was performed using an unpaired t-test: n.s. not significant, * p ≤ 0.05, ** p ≤ 0.01, *** p ≤ 0.001, **** p ≤ 0.0001.

### Genetic perturbation of ETC subunits rewires LGMN regulation *in vivo*

To test whether biasing cells towards an OXPHOS or glycolytic state *in vivo* tips the directional regulation of immune proteins, we used two KO mouse models lacking unrelated ETC-interacting factors (Fig. 4A). First, we used a *Ndufa4l2* KO line. *Ndufa4l2* is a HIF-1α induced ETC-interacting micropeptide, conserved in vertebrates, involved in mitochondrial stress responses (*22, 37*), and has viral homologs. *Ndufa4l2* displays pro-glycolytic and ETC Complex I inhibitory activity, *in vitro* (*37, 38*). We hypothesized that *Ndufa4l2* KO would bias mice towards the galactose/OXPHOS program during an immune response, manifesting as decreased LGMN in *Ndufa4l2^-/-^*mice relative to WT. To trigger an immune response, we used the endotoxin shock model, a well-appreciated method to promote mitochondrial dysfunction, a switch to glycolysis (*15*), and activate the HIF-1α response (*39*).

**Fig. 4.**
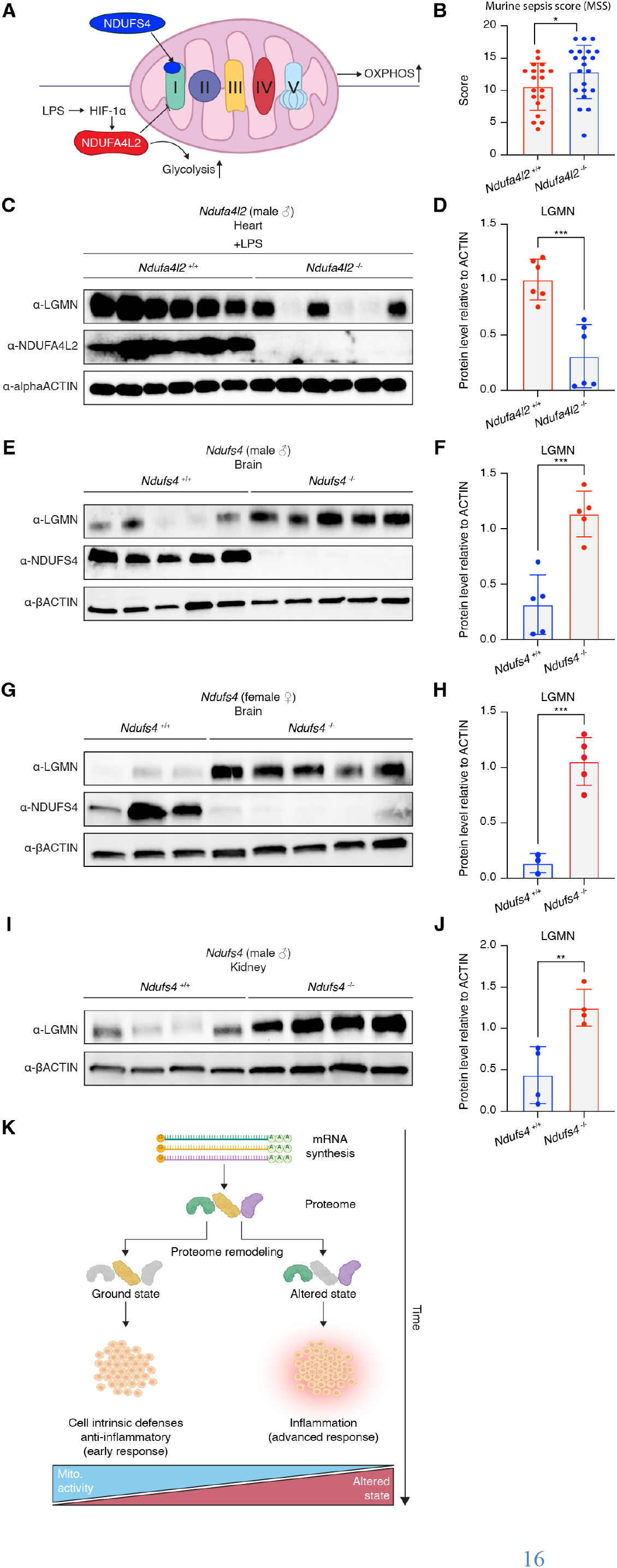
Genetic perturbation of ETC factors dysregulates *in vivo* expression of LGMN, an advanced immune response protein. (**A**) Schematic displaying the hypothesis for mice lacking NDUFA4L2, a pro-glycolytic and ETC Complex I inhibitor, following LPS-injection and a mouse model of mitochondrial disease that lacks NDUFS4, a core ETC Complex I subunit. (**B**) Murine sepsis scoring of 4 hours post-LPS injection. (**C, D**) Western blot and quantification of LGMN in heart lysates from *Ndufa4l2*^+/+^ and *Ndufa4l2*^-/-^ mice 4 hours post-LPS injection (i.p.). (**E, F**) Western blot and quantification of LGMN using brain lysates from untreated male *Ndufs4*^+/+^ and *Ndufs4*^-/-^. (**G, H**) Western blot and quantification of LGMN using brain lysates from untreated female *Ndufs4*^+/+^ and *Ndufs4*^-/-^. (**I, J**) Western blot and quantification of LGMN using kidney lysates from untreated male *Ndufs4*^+/+^ and *Ndufs4*^-/-^. (**K**) Model of mitochondria-mediated proteome remodeling. Statistical analysis was performed using an unpaired t-test: n.s. not significant, * p ≤ 0.05, ** p ≤ 0.01, *** p ≤ 0.001, **** p ≤ 0.0001.

*Ndufa4l2^-/-^* mice displayed no overt defects under homeostatic conditions. However, *Ndufa4l2^-/-^* mice manifested worse outcomes when injected intraperitoneally with LPS as evidenced by a more severe clinical sepsis score relative to WT littermate controls (Fig.4B, p < 0.05). Sepsis scores track appearance, consciousness, activity, response to stimulus, respiration rate, and respiration quality (*40*). Looking in the heart, a metabolically demanding organ affected by sepsis, LPS-injected *Ndufa4l2^-/-^* mice exhibited decreased LGMN protein levels relative to controls (Fig. 4, C and D, p < 0.001). These data suggest that an inhibitor of the ETC, which is pro-glycolytic, is necessary to upregulate LGMN - a marker of the “glucose” program - during an *in vivo* immune response.

Next, we tested whether ETC disruption was sufficient to induce expression of LGMN using untreated *Ndufs4^-/-^* mice. *Ndufs4* is a conserved ETC subunit, and KO mice display OXPHOS defects under homeostatic conditions (*41*). *Ndufs4^-/-^*mice are the quintessential model for mitochondrial disease and display symptoms similar to the neurological disorder Leigh syndrome including neuroinflammation, ataxia, reduced size, and short lifespan. In brain, a major site of pathology in humans and these mice (*42*), we found marked upregulation of LGMN in both male (Fig. 4, E and F, p < 0.001) and female *Ndufs4^-/-^* mice (Fig. 4, G and H, p < 0.001) relative to controls. We next examined the kidneys, which are sensitive to metabolic alterations and display defects in this disease. Similar to brain, we found increased LGMN in *Ndufs4^-/-^* kidneys relative to littermates (Fig. 4, I and J, p < 0.01). These data suggest deletion of a core ETC factor in mice is sufficient to promote expression of LGMN, a marker of the “glucose” program.

### Conclusions

This study provides evidence that perturbation of mitochondrial activity temporally regulates an appreciable subset of immune proteins by stratifying these factors into two previously unrecognized expression classes. While studies of primary immune signaling outcomes have investigated transcriptome changes in detail, changes at the protein level and the impact of secondary cues have been less explored. Nevertheless, metabolism is a well-appreciated modifier of immunity; yet its effects remain incompletely understood (*13, 43*). Given mitochondrial dysfunction is linked to inflammation in both infection and diverse chronic diseases (*3*), potential parallels between the catalysts, effectors, and outcomes of the activities have been implied and are now emerging.

Our *in vitro* and *in vivo* data suggest that the mitochondrial activity imparts an evolutionary conserved logic to the host response independent of immune cues such as type I IFN, type II IFN, and LPS (Fig. 1 to 4). Although anecdotal examples of immune proteins regulated by metabolism exist (COX2) (*44*), the breadth of immune proteins regulated by rewiring mitochondrial activity was unexpected. Equally striking is that the direction of regulation mirrors the progression of metabolic reprogramming during infection. Specifically, metabolism frequently shifts from the ground state of OXPHOS to other metabolic states like glycolysis, accompanied by the progression of immunity from cell autonomous defenses to later extrinsic responses like inflammation. For example, galactose media, which promotes OXPHOS, favors expression of host proteins with established roles in cell-intrinsic defense, negative regulators of inflammation, and even several viral entry factors. Given OXPHOS is common cellular state that viruses initially encounter, viruses would be predicted to evolve and co-opt factors expressed under these conditions. Likewise, the earliest host defense factors in the infected cell, along with anti-inflammatory molecules that inhibit advanced immune responses, would be expected to be stably expressed under OXPHOS. Moreover, mitochondrial perturbation seems to cue a transition in the proteome from cell autonomous defense factors to regulators of inflammation and professional immune cells. Proteins upregulated by glucose and ETC-perturbation include IRF1, LGMN, and enzymes COX2 and AKR1C1, which generate prostaglandins and inactivate progesterones, respectively. LGMN is interesting because it is linked to pathological processing of Tau, TDP-43, and α-synuclein, which are dysregulated in neurological diseases that display mitochondrial dysfunction (*45–47*).

Based on these data, we propose a model where the core metabolic state of OXPHOS promotes expression of proteins involved early in the response to pathogens whereas sustained perturbations to mitochondrial activity foster expression of downstream factors that are essential in processes like inflammation (Fig.4K). How mitochondrial activity regulates these proteins is unclear. However, it may be due to either translational control, protein degradation similar to selective IRF1 degradation in galactose media (*21*), or both. Interestingly, mTOR inhibitors (*48, 49*) regulate IFITM3 protein levels and antiviral activity. Future studies will inform how mitochondrial activity is sensed to directionally regulate early and late host response proteins. As this regulation is active in diverse cell types and pathogens represent a major selective force over evolutionary time, it may be an ancestral host defense response linked to mitochondria. Given the temporal contribution, it is tempting to speculate that sensing of mitochondrial dysfunction is evolutionary mismatched in metabolic diseases where OXPHOS perturbation is being sensed as an advanced infection.

## Acknowledgments

We express our gratitude to members of the Hancks Lab and Drs. Robert Orchard, Daniel Propheter, and Andrew Sandstrom for their feedback on the project and manuscript. We also express gratitude towards the UT Southwestern Proteomics core for help with data generation.

## Funding

National Institutes of General Medical Sciences R35GM142689, 1R35GM162011 (DCH)

Cancer Prevention Research Institute of Texas RR 170047 (DCH)

National Institutes of Health grant T32 AI007520 (JA)

National Institutes of Health grant 2T32AI005284-41A1 (TC)

## Author contributions

Conceptualization: JA, DCH

Methodology: JA, SR, SB, NH, TC, KKC, JL, AL, DCH

Investigation: JA, SR, SB, NH, TC, KKC, JL, AL, DCH

Visualization: JA, SR, SB, NH, TC, KKC, DCH

Funding acquisition: JA, TC, DCH

Project administration: JA, TC, DCH

Supervision: DCH

Writing – original draft: JA, DCH

Writing – review & editing: JA, DCH

## Competing interests

Authors declare that they have no competing interests.

## Data, code, and materials availability

All data are available in the main text or the supplementary materials. The mass spectrometry proteomics data have been deposited to the ProteomeXchange Consortium via the MassIVE partner repository with the dataset identifier PXD083611 and MassIVE ID MSV000103102. The data can be accessed at ftp://massive-ftp.ucsd.edu/v14/MSV000103102/. RNA-seq data are publicly available in the NCBI GEO database under accession GSE226242.

## Notes

### Competing Interest Statement

The authors have declared no competing interest.

